# Reconceptualizing beta diversity: a hypervolume geometric approach

**DOI:** 10.1101/2022.11.27.518099

**Authors:** Chuliang Song, Muyang Lu, Joseph R. Bennett, Benjamin Gilbert, Marie-Josée Fortin, Andrew Gonzalez

## Abstract

Beta diversity—the variation among community compositions in a region—is a fundamental measure of biodiversity. Despite a diverse set of measures to quantify beta diversity, most measures have posited that beta diversity is maximized when each community has a single distinct species. However, this assumption overlooks the ecological significance of species interactions and non-additivity in ecological systems, where the function and behaviour of species depend on other species in a community. Here, we introduce a geometric approach to measure beta diversity as the hypervolume of the geometric embedding of a metacommunity. This approach explicitly accounts for non-additivity and captures the idea that introducing a unique, species-rich community composition to a metacommunity increases beta diversity. We show that our hypervolume measure is closely linked to and naturally extends previous information- and variation-based measures while providing a unifying geometric framework for widely adopted extensions of beta diversity. Applying our geometric measures to empirical data, we address two long-standing questions in beta diversity research—the latitudinal pattern of beta diversity and the effect of sampling effort—and present novel ecological insights that were previously obscured by the limitations of traditional approaches. In sum, our geometric approach reconceptualizes beta diversity, offering an alternative and complementary perspective to previous measures, with immediate applicability to existing data.

## 1 Introduction

Beta diversity is one of the most important measures of biodiversity (Anderson *et al*., 2011; Mittelbach & McGill, 2019). In essence, beta diversity aims to measure the diversity of between-community components, or the number of effective communities. It serves as a bridge connecting ecological phenomena from local to regional scales. To maintain consistency with existing literature, we use the terms ‘community’ and ‘site’ interchangeably throughout this paper. Unfortunately, it remains one of the most debated concepts in biodiversity research. Since the concept was conceived in the mid-20th Century (Whittaker, 1960, 1972), researchers have come up with a long list of measures (reviewed in Anderson *et al*. 2011). Some recent notable measures include Hill numbers (Jost, 2007; Ohlmann *et al*., 2019), *β*-deviation (Kraft *et al*., 2011; Xing & He, 2021), turnover-nestedness decomposition (Baselga, 2012; Legendre, 2014), and variance of community composition matrix (Legendre & De Cáceres, 2013). Importantly, although these measures focus on different aspects of beta diversity, most of them obey the same set of mathematical axioms (Legendre & De Cáceres, 2013). However, some of these axioms may not align with the key ecological intuition on what beta diversity should be.

The key discrepancy rests on what it means for a community to be different from other communities. To explain the problem in a nutshell, let us consider two metacommunities (labeled as I and II; Figure 1), both with 2 species (labeled as *A* and *B*), and either 2 or 3 local communities (labeled as 1-3). For simplicity, we use the Whittaker’s multiplicative measure of beta diversity (*β* = *γ/α*) to represent the traditional measures. Metacommunity I (Figure 1A) has one community with only species *A* and another with only *B*, leading to a beta diversity of 2 (as *γ* = 2 and *α* = 1). In contrast, metacommunity II (Figure 1A) adds a third community containing both species *A* and *B*, resulting in a beta diversity of 1.5 (*γ* = 2 and *α* = 4*/*3). Thus, the traditional measures argue that metacommunity I has a larger beta diversity than metacommunity II (Figure 1B). Note that this is not a special property of the Whittaker’s multiplicative measure but satisfied by almost all measures (reviewed in Legendre & De Cáceres 2013). This result means that by adding a unique, distinctive community ({*A, B*}) to existing communities, the beta diversity of the metacommunity would decrease.

**Figure 1:**
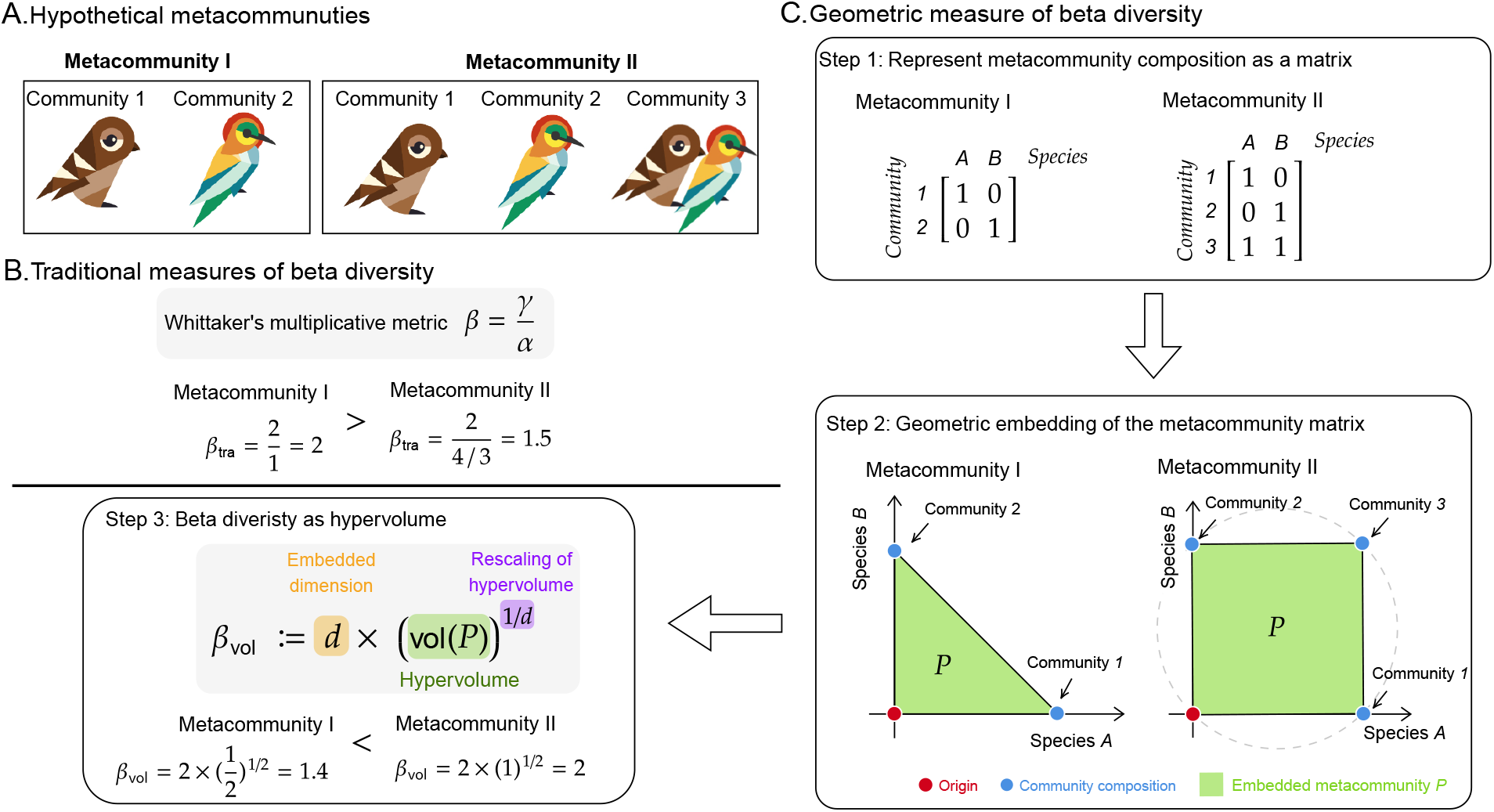
Illustration of the geometric approach to beta diversity. Panel (A) shows two hypothetical metacommunities (labeled as I-II) with 2 species (labeled as *A* and *B*) and up to 3 local communities (labeled as 1-3). The two communities of metacommunity I consist of species *A* only and species *B* only, respectively. The three communities of metacommunity II consist of species *A* only, species *B* only, and species *A, B* together, respectively. Panel (B) shows that the traditional measures of beta diversity asserts that metacommunity I has a *higher* beta diversity than metacommunity II. We show only the case for Whittaker’s multiplicative metric, but the qualitative order would be the same for all traditional measures of beta diversity. Panel (C) illustrates the key steps of our geometric measure. The first step is to turn the metacommunities in Panel (A) into the equivalent matrix representations. This matrix is known as community composition matrix, where the rows represent communities, the columns represent species, and elements represent species presence/absence. More generally, the elements can be any measure of species importance, such as abundance or biomass. The second step is a geometric embedding of the metacommunity. Here, as we have a smaller number of species than the number of communities, the number of species determines the dimension and axis of the embedded space (2-dimensional space), while the communities determined the embedded points (blue points). Note that the origin (red point) is automatically embedded because adding an empty community should not affect beta diversity. The metacommunity is now realized as the spanned geometric object (green area; denoted as *P*) by all the embedded point and the origin. The third step is to measure the beta diversity as the normalized hypervolume of the geometric object: *β*_vol_ = *d* × (vol(*P*))^1*/d*^, where *d* is the embedded dimension (2 here as there are 2 species). Within this measure, metacommunity I has a *lower* beta diversity than metacommunity II, opposite to the traditional measures.

Fundamentally, the traditional measures assume that the community {*A, B*} is *not different*, or even redundant, from the community {*A*} and community {*B*}. This assumption ignores, however, the importance of species interactions and non-additivity in ecological systems Extensive research on biodiversity-ecosystem function (Tilman *et al*., 2014; Gonzalez *et al*., 2020), trait-mediated indirect interactions (Werner & Peacor, 2003; Ohgushi *et al*., 2012), higher-order interactions (Kelsic *et al*., 2015; Majer *et al*., 2024), and foodweb stabilization through weak interactions (McCann *et al*., 1998; Neutel *et al*., 2002), has shown that *a community is more than the sum of its parts*: a community with multiple interacting species is ecologically different from a set of isolated communities each containing a single species. Traditional beta diversity measures fail to capture this critical aspect of ecological systems. In the example from Figure 1, the community {*A, B*} behaves differently, both dynamically (Levine *et al*., 2017; Angulo *et al*., 2021) and functionally (Maron *et al*., 2018; van der Plas, 2019), from the communities with only {*A*} or {*B*}. Consequently, we argue that community {*A, B*} should be considered *different* from the communities with only {*A*} or {*B*}. To generalize, the introduction of a unique, distinctive community composition to a metacommunity should increase the diversity within it. Following this rationale, opposite to traditional measures, we should assign a higher beta diversity for metacommunity II than metacommunity I (Figure 1C).

To resolve this discrepancy, we introduce a new measure of beta diversity using an intuitive and visual geometric approach. The key idea of this geometric approach is to view the metacommunity as a geometric object occupying hyperspace, and then quantify its beta diversity as the hypervolume of the geometric object. Firstly, we illustrate the key ideas with simple examples, and provide a generalization to metacommunities with arbitrary structure. Then, armed with this geometric perspective, we provide a unified treatment of common variants beyond basic beta diversity: duplications in presence/absence data, temporal changes, community/species-specific contribution, turnover-nestedness decomposition, and accounting for species similarity and functional complementarity. In contrast, traditional approaches require different formalisms to deal with these variants. We then show this geometric approach is linked to and naturally extends classic measures of beta diversity based on generalized covariance and information theory. Lastly, we apply our hypervolume measure of beta diversity to empirical datasets, including the trend of beta diversity along longitude and the sampling efforts.

## 2 Geometry of beta diversity

Most definitions of beta diversity stem from algebraic manipulations of metacommunity properties (Anderson *et al*., 2011). Here, we provide an alternative geometric approach. This approach is grounded on the idea of embedding an arbitrary metacommunity as a hyper-dimensional geometric object. We will show that this geometric shape of metacommunity provides a unifying bridge to various definitions of beta diversity.

### 2.1 Illustration of the basic idea

To illustrate the basic idea, let us consider again the hypothetical examples of metacommunities in Figure 1. Recall that in metacommunity II, community 1 only has species *A*, community 2 only has species *B*, and community 3 has both species *A* and *B* (middle panel in Fig 1A). We can represent the metacommunity in a matrix form (Figure 1B):

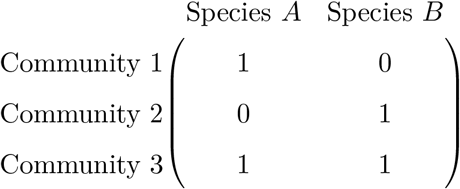

where the columns denote species, the rows denote the communities, and the elements denote whether the given species is present in the given community (1 for presence and 0 for absence). We call this matrix form the *metacommunity matrix*. Note this matrix form is also known as community matrix in the literature (Legendre & De Cáceres, 2013).

The crux of our new definition of beta diversity is to interpret this matrix as points in a hyper-dimensional space. In this example, the space is 2-dimensional (each species as an axis) and we have three points (rows in the matrix: (1, 0), (0, 1), and (1, 1)). The middle panel (step 2) of Figure 1C illustrates the geometric embedding of the matrix. Beta diversity is related to the volume spanned by these points together with the origin. The ecological rationale to add the origin is known as double-zero asymmetry (appendix S3 of Legendre & De Cáceres 2013): beta diversity should not change when we “add” a ghost species that does not exist in any of the communities (which is the origin in the space), because such a ghost species is not interpretable (Whittaker, 1972). Thus, the origin must be included for ecological consistency of beta diversity.

With this geometric embedding (Figure 1C), we can see that metacommunity II, which has three distinct community compositions, has a volume of 1 as a square with side length 1. In comparison, metacommunity I, which has two distinct community compositions, has a volume of 0.5. Thus, this hypervolume approach naturally resolves the discrepancy regarding when beta diversity is maximized: more distinct compositions correspond to more unique points in the hyper-dimension, which leads to greater hypervolumes. Note that the hypervolume would be minimized (= 0) if all communities have identical compositions, which align with the intuition of beta diversity with non-additivity.

To make the hypervolume measure representing an effective number of communities with 2 species (Jost, 2007), we define beta diversity *β*_vol_ in this example as the rescaled volume of the raw volume 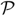:

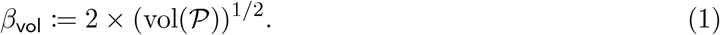

With this definition, the metacommunity I has a beta diversity of 1.4, while the metacommunity II has a beta diversity of 2 (the highest possible beta diversity).

### 2.2 Generalization to arbitrary metacommunity

We can generalize the simple cases above to complex metacommunities. For a general metacommunity with *N* local communities and *γ* species, we can represent it using a general metacommunity matrix **Z**, where each row represents a local community and each column represents a species:

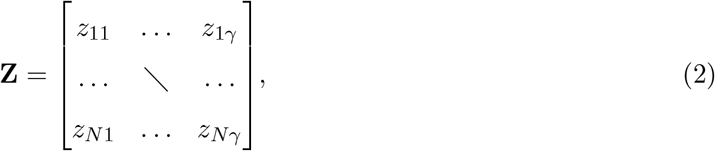

where the element *z*_*ij*_ represents an ecological measure of the importance of species *j* in local community *i*. This measure can be any measured value, such as presence (1 if present, 0 if absent), abundance (number of individuals), or biomass (total mass of the species in the community). A caveat, though, is that *z*_*ij*_ need to be appropriately scaled to make it fully comparable across metacommunities and to avoid the issue of “points-in-the-middle” problem (Legendre & De Cáceres, 2013).

We need to identify the constraint on beta diversity: the gamma diversity (*γ*), or the number of communities (*N*). The smaller of these two values acts as the constraint, determining the dimension of the embedded space, while the larger value represents the number of embedded points. This identification ensures that beta diversity is well-defined for all metacommunities. The motivation behind the constraint is to align with the concept of maximum effective communities in traditional measures. For the case when there are more communities than species, the Whittaker’s multiplicative measure asserts that the maximum beta diversity cannot exceed *γ* (because the minimum alpha diversity is 1 as every community has at least one species). Conversely, for the case when there are more species than communities, the Whittaker’s multiplicative measure asserts that the maximum beta diversity cannot exceed *N* (because the minimum alpha diversity is *γ/N* as every species would occur at least once in some community), which is the constraint.

Formally, the expanded convex hull 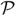 of the geometrically embedded points (representing communities) in the *d* = min(*γ, N*) -dimensional space is

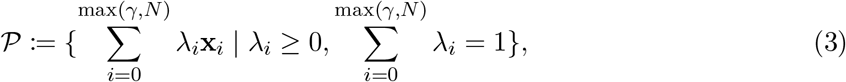

where *x*_0_ corresponds to the origin, **x**_*i*_ corresponds to either the *i*-th column or row of the metacommunity matrix (depending on which is the constraint), and *λ*_*i*_ corresponds to the weights associated with each point (community).

Following the definition above, our measure of beta diversity *β*_vol_ (the underscript highlights the use of hypervolume) is defined as the rescaled hypervolume of the convex hull 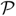:

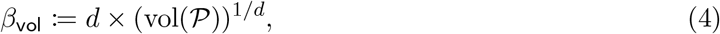

where *d* = min(*γ, N*) is the constraint and (vol(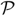))^1*/d*^ is the normalized hypervolume. *β*_vol_ is interpreted as the number of effective communities, which ranges from 0 to *d*.

This rescaling of raw hypervolume in Eqn. 4 is fundamental for its interpretation as beta diversity. A heuristic argument is that, with *γ* species, the hypervolume beta diversity should range from 0 (achieved with only 1 unique community composition) to *γ* (achieved with (2^*γ*^ *−* 1) distinct community compositions). This range of beta diversity is based on the argument that the effective number is mostly ecologically intuitive (Jost, 2007). To get rid of the effects of the exponential increase of distinct community compositions, we need to take the *γ−*th root of the hypervolume. Of course, further rescaling of Eqn. 4 is possible depending on different ecological rationales (e.g., beta diversity should range from 0 to 1).

To validate the heuristic argument behind the rescaling of the hypervolume (Eqn. 4), we compute all possible beta diversities for metacommunities with three species (i.e., *γ* = 3). The maximum beta diversity (*β*_vol_ = 3) is achieved with (2^*γ*^ *−* 1) = 7 distinct community compositions, while the minimum beta diversity is achieved with only 1 distinct community composition. Figure 2 shows the rescaled hypervolume has positive saturating association to the number of unique community compositions in a metacommunity. Appendix B shows the linear scaling persists for higher gamma diversity.

**Figure 2:**
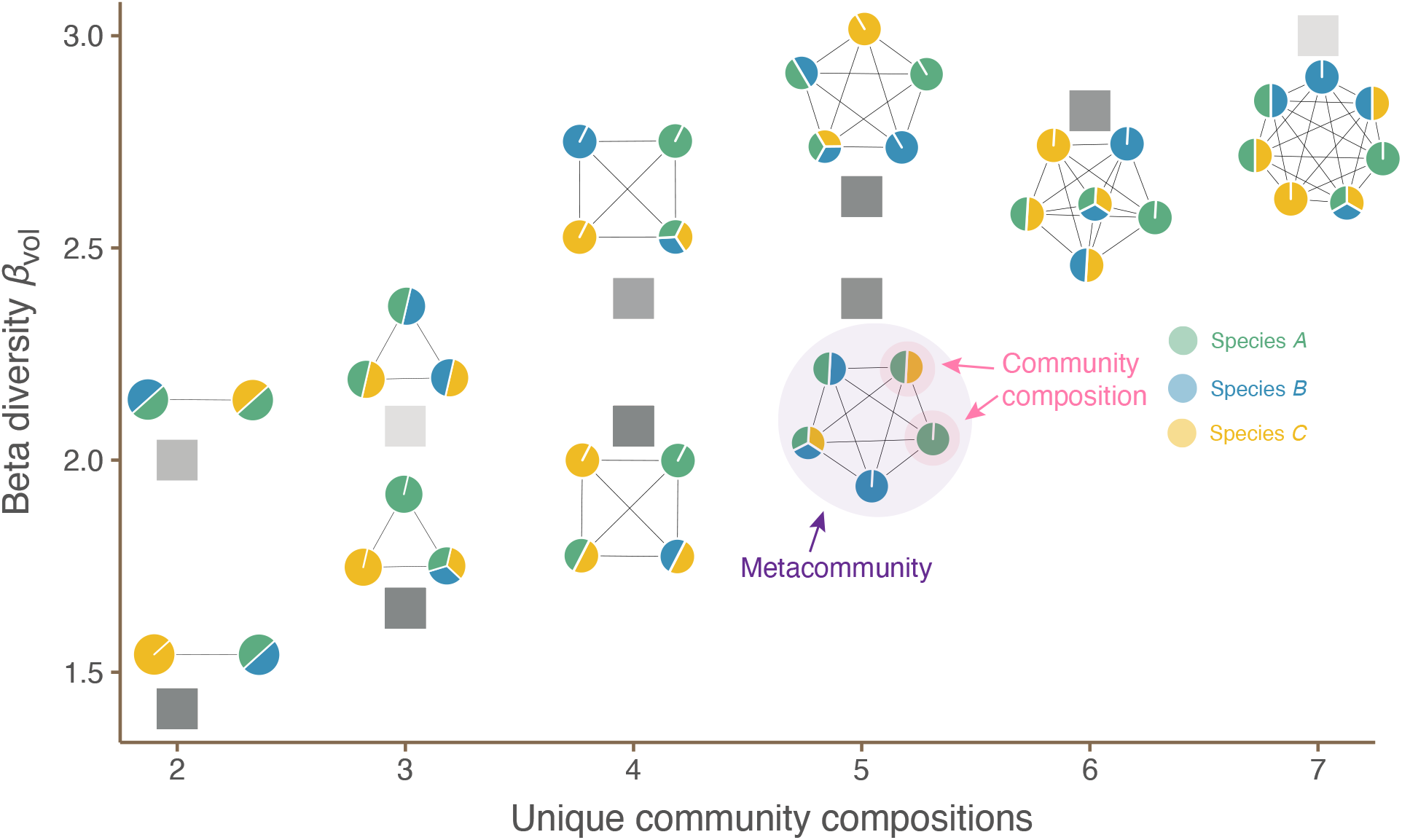
Possibilities of beta diversities for metacommunities with gamma diversity = 3. For simplicity, here we only consider metacommunities with information on species presence or absence. The horizontal axis shows the number of unique community compositions in the metacommunity, while the vertical axis shows the hypervolume beta diversity (*β*_vol_) defined in Eqn. (4). For clarity of presentation, we only show metacommunities with nontrivial beta diversity (i.e., *β*_vol_ *>* 0). The transparency of the square denotes the number of distinct metacommunities that have identical beta diversity with the same number of unique community compositions, with more solid squares indicating more metacommunities. For each square, we illustrate one example of metacommunity. In each metacommunity, the communities are represented as nodes, and the colors of the nodes represent community compositions (green, blue, and yellow for species *A, B, C*, respectively). Beta diversity *β*_vol_ increases with the number of unique community compositions in a linear trend with notable variations. These variations are due to different levels of similarities in species compositions.

In addition to the positive trend, values of beta diversity have notable variations within the same number of unique community compositions. Thus, even though the number of unique community compositions is the key determinant of beta diversity, how distinct the unique community compositions are is also another determinant. For example, a community with species composition {*A, B*} is more distinct from a community with composition {*C*} than a community with composition {*A*}. In other words, *β*_vol_ does not treat all unique compositions equally; instead, it puts higher weights to more distinct community compositions.

## 3 A unified framework of beta diversity

In the previous section, we have introduced a geometric approach to define beta diversity through a geometric embedding of a metacommunity (Eqn. 4). We have so far only focused on the most basic case of beta diversity. Many important extensions of beta diversity have been proposed through the study of beta diversity, such as temporal dimension (De Cáceres *et al*., 2019) and accounting for species similarity (Leinster & Cobbold, 2012). Despite their importance, these extensions require different methodologies. With the flexibility empowered with geometry, here we provide a unified treatment to these extensions in beta diversity theory.

### 3.1 Duplications in presence/absence data as weighted embedding

Information on species presence or absence is often the only available data in empirical metacommunities. Mathematically, this means *z*_*i*_ = 1 or 0 in the metacommunity matrix (Eqn. 2). A common issue with these data is the duplication of identical community compositions. However, the definition of *β*_vol_ (Eqn. 4) in the previous section does not take this into account because communities with duplicated compositions all map to the same embedded point.

To account for this, we provide a simple modification to account for duplicated community compositions through weighted embedding. We compress all communities with duplicated compositions into one community and then assign the frequency of identical communities as its weight. Figure 3 illustrates an example of metacommunity with 2 species and 6 communities.

**Figure 3:**
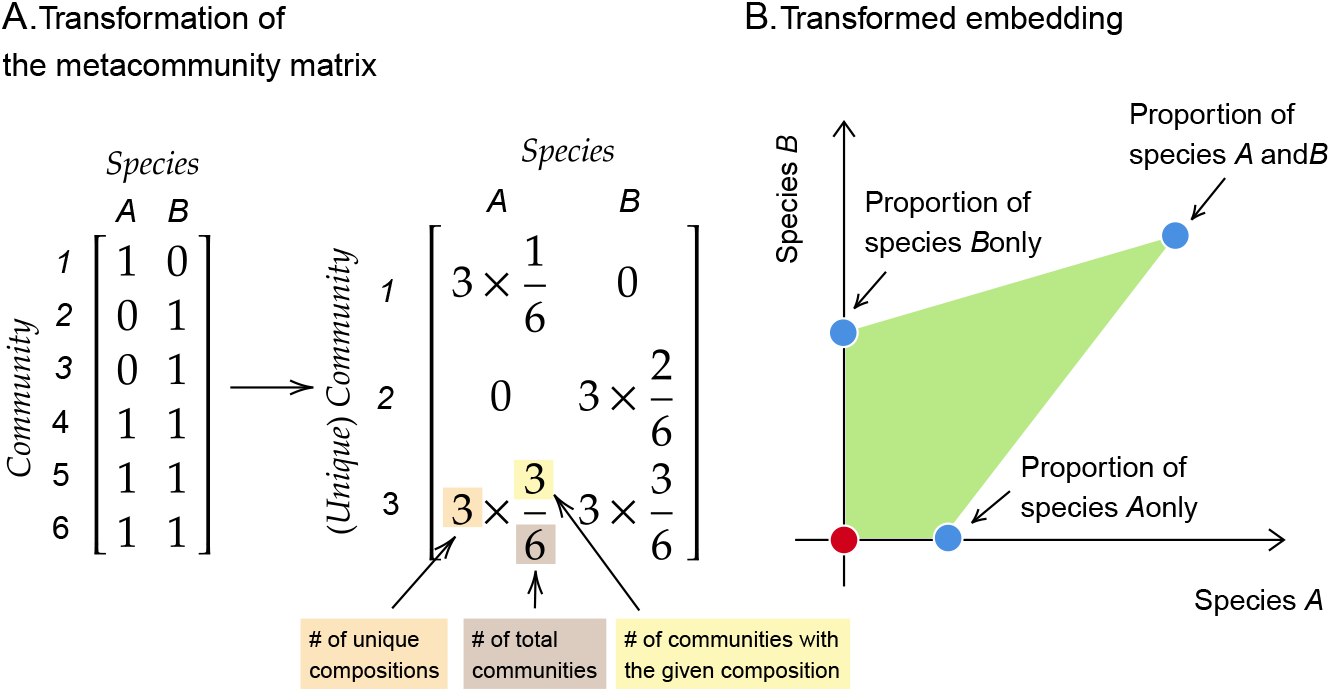
Weighted geometric embedding of metacommunity with presence/absence data. Panel (A) shows an example of metacommunity with 2 species and 6 communities. The left matrix is the original metacommunity matrix, while the right one is the transformed meta-community matrix. This weighted transformation is given by (# of unique compositions) × 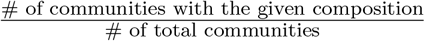. This embedding scheme ensures that metacommunities without duplications would be identical after the transformation. Panel (B) shows the transformed embedding of the metacommunity.

Formally, suppose we have *N* local communities and *S* species in a metacommunity where the number of species is the constraint (i.e., *S < N*). The same modification can be applied when the number of communities is the constraint. The species composition of the *i*-th community is **x**_*i*_ := {*z*_*ij*_}. Then suppose among the *N* communities, we have only *m* effectively unique communities **y**_*k*_ (*k* = 1, …, *m*), where each unique community **y**_*k*_ appears *n*_*k*_ times. Then

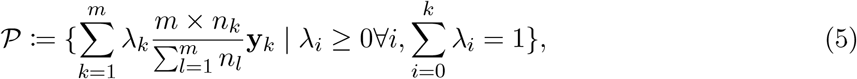

where 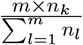 provides the weighted embedding of *y*_*k*_ and *λ*_*k*_ generates the convex hull. The weight would be 1 if all communities have distinct compositions (i.e., *n*_*k*_ = 1, ∀*k*). Note that it is straight-forward to apply this modification to weighted embedding to other types of measures of species importance, although in empirical data, the modification is unlikely to be needed. For example, it is unlikely that two communities would have identical abundances for all their constituent species.

As an application, we can ask the following question: for a metacommunity with 2 species (labeled as *A* and *B*), what is the proportion of communities with compositions {*A*}, {*B*} and {*A, B*} that maximize the beta diversity? Our metric reaches a maximum when 1*/*4 communities have {*A*}, 1*/*4 communities have {*B*}, and the other 1*/*2 communities have {*A, B*} (Appendix A).

### 3.2 Temporal turnover of beta diversity as hypervolume overlap

Beta diversity *per se* is a measure on the spatial scale. To fully understand biodiversity changes, we need to study how beta diversity changes over time and over different temporal scales (Gonzalez *et al*., 2020). One approach is to directly compare beta diversity values at two distinct time points, revealing the magnitude of change in among-site differences. While straightforward, this method overlooks the crucial aspect of turnover—the extent to which community compositions synchronize across the entire metacommunity. To capture this information, De Cáceres *et al*. (2019) proposed a method based on trajectory distances. Here, we measure the temporal change using the overlap between two geometric embeddings of metacommunities.

To illustrate the idea, let us consider a metacommunity with 2 communities and 2 species. At time (*t*), community 1 has only species *B* while community 2 has both species *A* and *B* (Figure 4A). Then at time (*t* + 1), community 1 still has species *B* while community 2 now only has species *A* (Figure 4B). To compute the hypervolume overlap, we need to assign the orientation of the geometric embedding. This orientation specifies the direction of synchronization in the metacommunity. Note that the specific choice of orientation does not matter as long as it is fixed throughout time. Without loss of generality, we choose the orientation from origin to community 1 to community 2. Once we assign the orientation, the hypervolume would have signs, which means the hypervolume can be negative. From time (*t*) to (*t* + 1), the orientations of the geometric embeddings do not change (both are clockwise). The hypervolume overlap is simply the overlap between two positive hypervolumes, which equals to 1*/*4.

**Figure 4:**
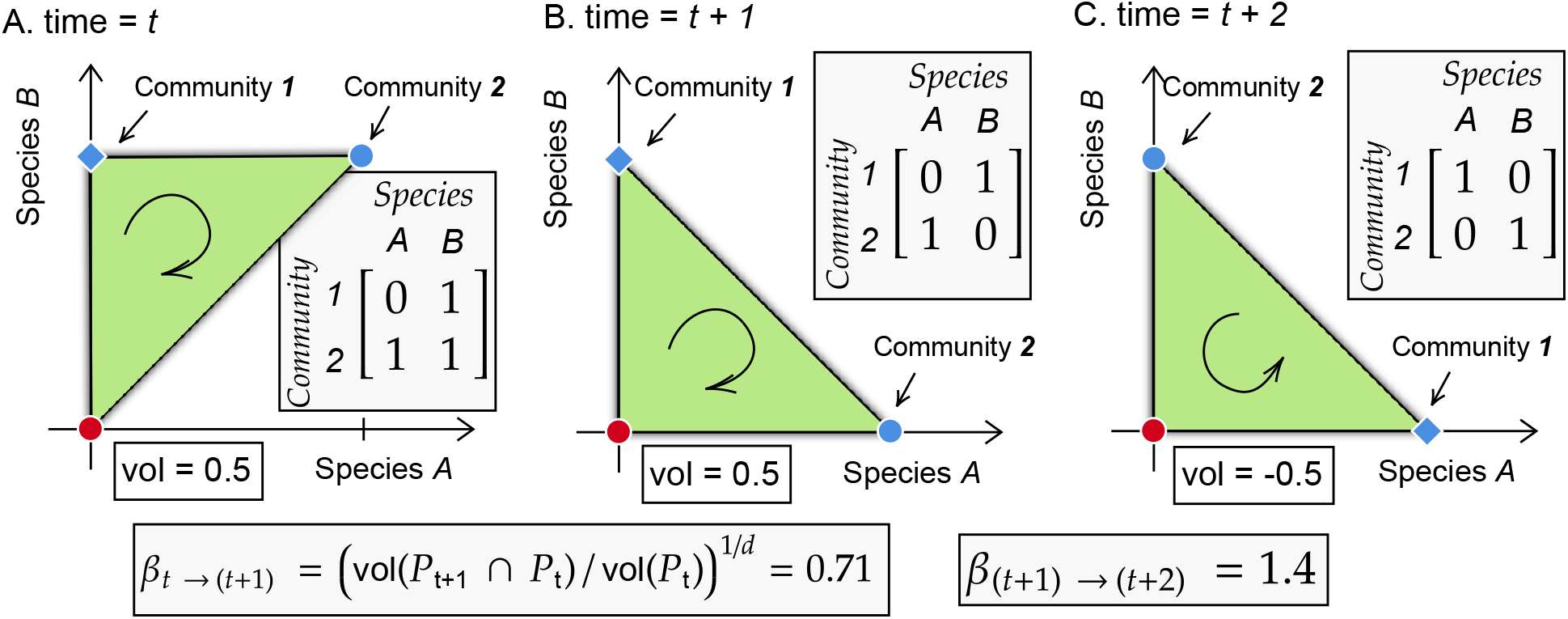
Measure temporal changes of beta diversity using (oriented) hypervolume overlap. We assign an orientation of hypervolume from origin to community 1 to community 2. The ecological interpretation of the orientation is the direction of synchronization in the metacommunity. Panels (A-C) represent metacommunity at time (*t*) to time (*t* + 2), respectively. From time (*t*) to time (*t* + 1), community 1 is unchanged while community 2 loses species *B*. The changes in community compositions are *asynchronized*, which are reflected in the identical orientations of their hyper-volumes. In contrast, from time (*t* + 1) to time (*t* + 2), community 1 and community 2 switch their community compositions. The changes in community compositions are *synchronized*, which are reflected in the opposite orientations of their hypervolumes. With the definition of temporal change (Eqn. 6), beta diversity changes by 0.71 from time (*t*) to time (*t* + 1), while it changes by 1.4 from time (*t* + 1) to time (*t* + 2).

Let us consider another example. Suppose at time (*t* + 2), community composition switches from time (*t* + 1) (i.e, community 1 only has species *A*, while community 2 only has species *B*; Figure 4C). In this case, the orientations of the geometric embeddings are opposite (clockwise versus anti-clockwise). The overlap now needs to consider the signed difference, which equals to 0.5*−*(*−*0.5) = 1 (despite the seemingly identical shape).

Formally, we can define the changes of beta diversity from time (*t*) to time (*t* + 1) as

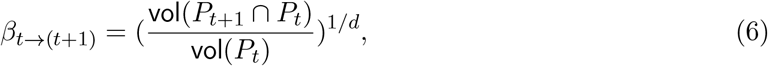

which measures the extent of synchronous or asynchronous changes in community composition in the entire metacommunity. If |*β*_*t*→(*t*+1)_ |*<* 1, then changes in community compositions are asynchronous or synchronous in the same direction. In contrast, if |*β*_*t→*(*t*+1)_ |*>* 1, then changes in community compositions are synchronous in the opposite direction.

Applying this definition (Eqn. 6) to the examples above, the change in beta diversity equals to 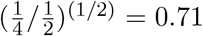 from time (*t*) to time (*t* + 1), while equals to 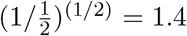 from time (*t* + 1) to time (*t* + 2). These results align with ecological intuition of ecological changes. From time (*t*) to time (*t* + 1), we see an asynchronous change in community compositions (community 1 remains fixed and only community 2 changes), which is reflected in a relatively smaller temporal change of beta diversity. In contrast, from time (*t* + 1) to time (*t* + 2), we see a synchronous change of community compositions (community 1 and community 2 switches composition), which is reflected in the relatively large temporal change of beta diversity.

### 3.3 Community/Species-specific contribution as hypervolume change

Communities do not contribute equally to biodiversity maintenance in a landscape. Thus, we need to disentangle the importance of community-specific contribution to beta diversity. Here, we measure the community-specific contribution using the relative change of hypervolumes.

From the perspective of our geometric approach, a given community contributes to the overall beta diversity through its embedded points. Thus, to evaluate its relative contribution, we can compare the overlap between the hypervolumes with and without this community. To illustrate, we use the metacommunity example in Figure 3. Figure 5A shows the original metacommunity matrix and its embedded geometric object. Figure 5B-D shows geometric objects without site 1-3, respectively. A key observation is the redundancy in beta diversity contributions from different communities. This is evident in Figure 5, where the sum of hypervolumes in B-D exceeds the original metacommunity’s hypervolume. To address this redundancy, we introduce a normalization step. Formally, the contribution of community *i* to beta diversity is

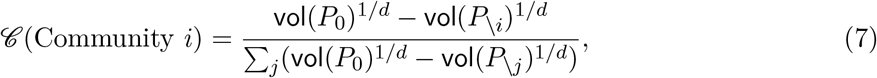

where *P*_0_ denotes the geometric object of the original metacommunity containing community *i, P*_*\ i*_ denotes the geometric object of the metacommunity without community *i*, and the summation index *j* runs through all communities. Applying Eqn. 7 to the above example (Fig. 5), we found community 1 contributes 0.16, community 2 contributes 0.37, while community 3 contributes 0.47.

**Figure 5:**
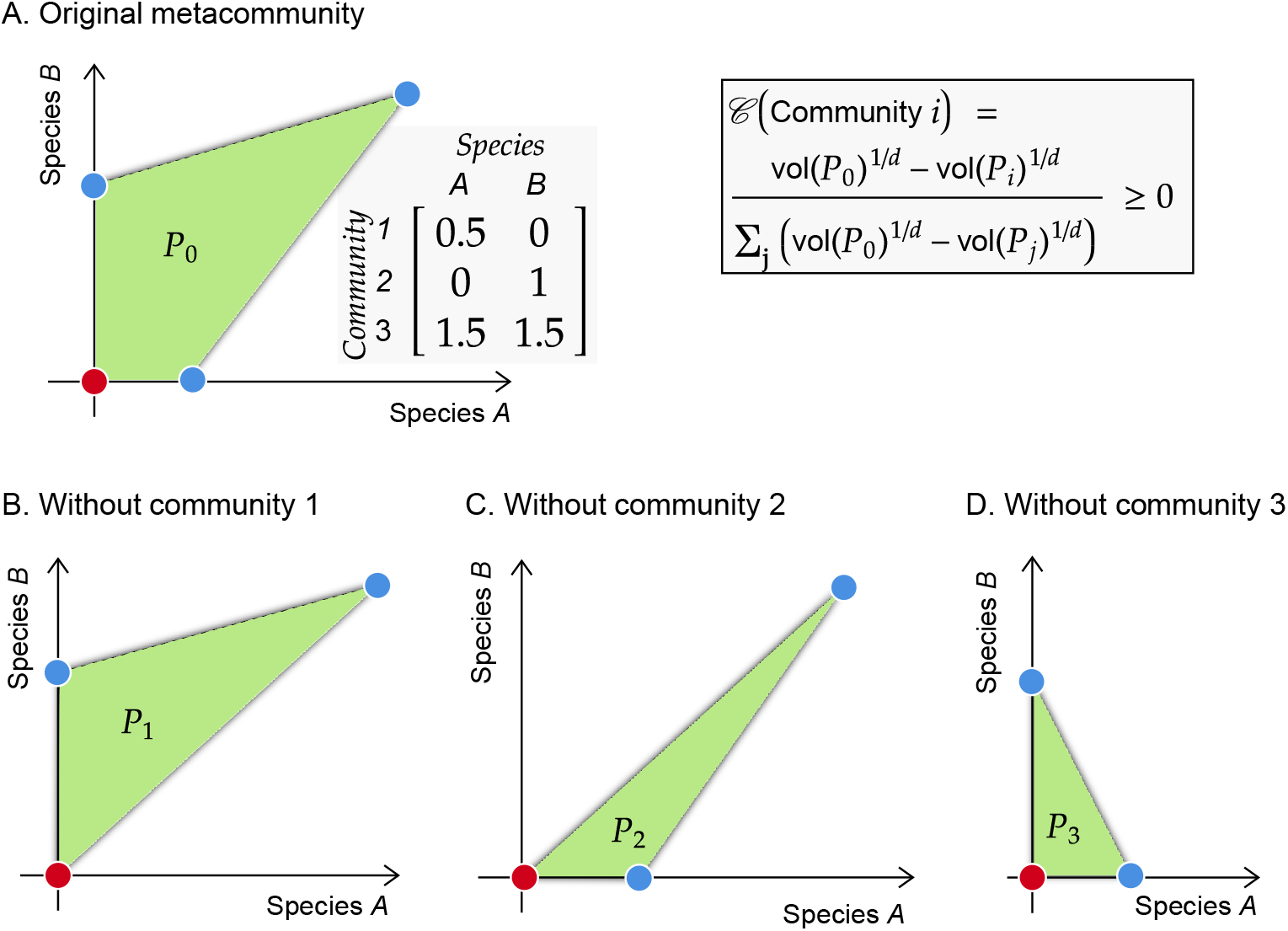
Disentangling site-specific contribution to beta diversity. Panel (A) shows an example metacommunity and its corresponding geometric embedding (the same as the example in Figure 3). Panels (B-D) show the metacommunities without community 1-3, respectively. The contribution of a community to the overall beta diversity is quantified as the normalized change in the hypervolumes. In this example, community 1 contributes 0.16, community 2 contributes 0.37, and community 3 contributes 0.47.

An important feature of our measure is that *all* communities have a non-negative contribution to beta diversity. This is because the hypervolume vol(*P*_0_) of the original metacommunity is always greater than or equal to that vol(*P*_*i*_) of the metacommunity without a community (under the assumption of not using the modified schemes on duplications (Eqn. 5) and no changes in *γ* diversity). In contrast, in the traditional formalization of beta diversity, a community may have a negative contribution (i.e., its presence decreases the beta diversity). For example, in the metacommunity II in Figure 1B, community 3 would have a negative contribution with the traditional formalization (e.g., it decreases beta diversity by 25% with Whittaker’s multiplicative measure). However, *ceteris paribus*, conservation management, in general, should *not* assign some community to be ‘negative’ for biodiversity (Hunter Jr & Gibbs, 2006). Thus, our framework is more appropriate to assess community contribution in conservation planning.

The above method parallelly applies to quantify species-specific contribution. A caveat is that we can only assess the contribution of either species or community (depending on which is the constraint on the embedded dimension), but not both.

### 3.4 Species similarity and functional complementarity as transformed embedding

Species are more similar to some species than others. To account for species similarity, we follow Leinster & Cobbold (2012) by introducing a *S* matrix where elements *s*_*ij*_ denote how similar species *i* is to species *j. s*_*ij*_ are scaled between 0 (totally dissimilar) to 1 (totally similar). For example, it can be a genetic or phenotypic (trait) similarity. Note that the *S* matrix is not required to be symmetric (i.e., *s*_*ij*_ *≠ s*_*ji*_).

From our geometric perspective, the *S* matrix corresponds to a linear transformation of the embedded geometric object. For simplicity, let us consider 2 species. Originally, (1, 0) denotes the presence of species *A* while (0, 1) denotes the presence of species *B*. The two axes are orthogonal. With the introduction of the *S* matrix, the presence of species *A* is now indicated as 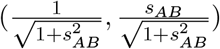 while the presence of species *B* is now indicated as 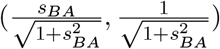. If all species are totally dissimilar, then the *S* matrix is an identity matrix. This corresponds to the same original axis (which is what we have been presenting so far; Figure 6A). For another example, if all species are similar, then the *S* matrix is a matrix with all 1s. This corresponds to all axes pointing to the exact same direction (1, 1) (Figure 6B). In this case, the hypervolume would always be 0. This agrees with ecological intuition, because the system effectively only has 1 species and there is no beta diversity. For a simple example, let us consider the *S* matrix 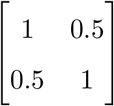. The hypervolume is now shrunk into a smaller region (Figure 6C).

**Figure 6:**
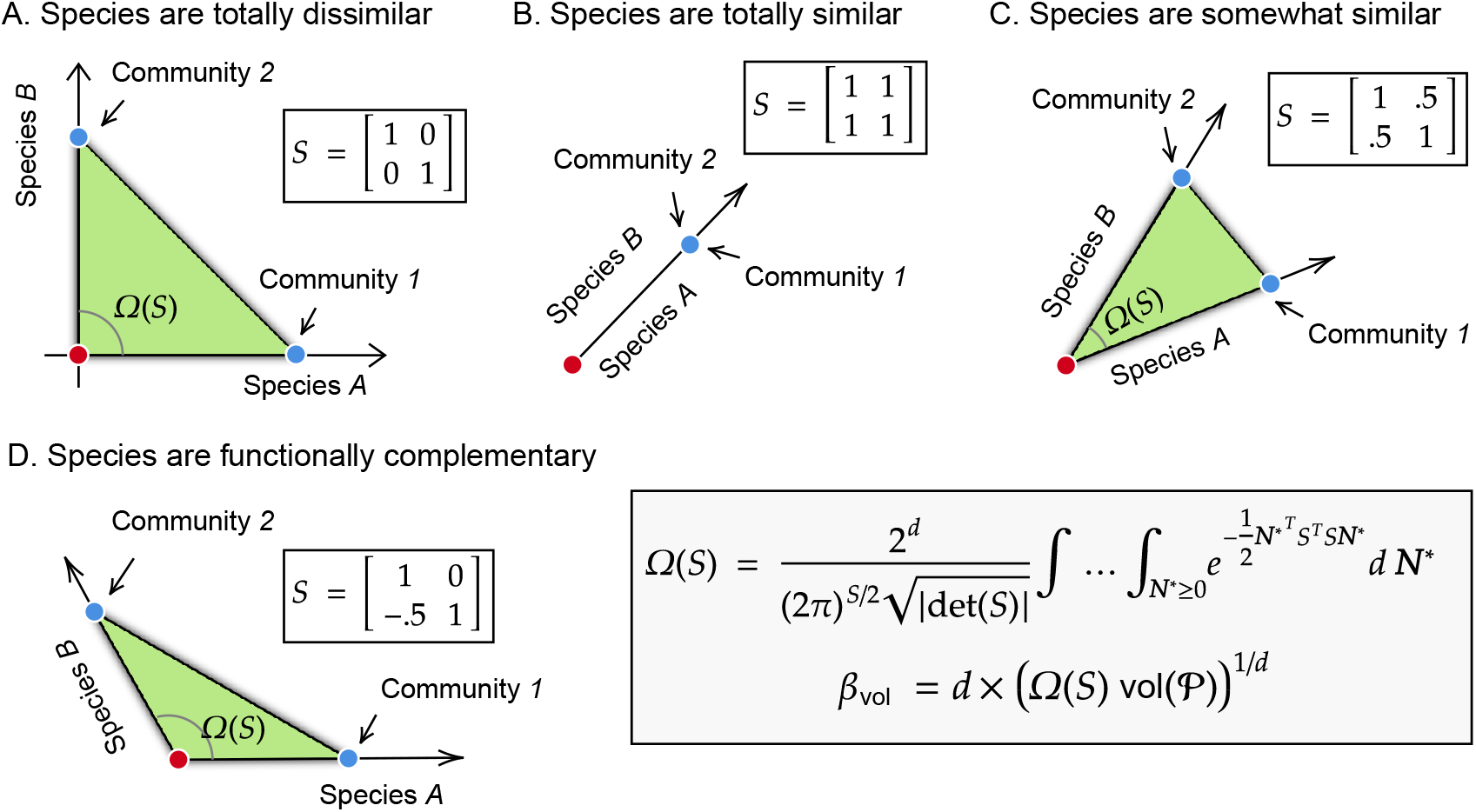
Accounting for species similarity and functional complementarity to quantify beta diversity. is equivalent to a coordinate transformation. All panels show the same metacommunity with different species’ similarity or functional complementarity. The original metacommunity has two communities where one community only has species *A* and the other community only has species *B* (the same as metacommunity II in Figure 1A). Panel (A) shows the case where species are totally dissimilar. The hypervolume and the corresponding beta diversity remain the same. Panel (B) shows the case where species are totally similar. The hypervolume shrinks to 0 and there is no beta diversity. Panel (C) shows the case where species are a bit similar. The hypervolume is larger than 0 but shrinks compared to the case where the totally dissimilar case. Panel (D) shows the case where species are functionally complementary. This is reflected in *S*_21_ *<* 0. The hypervolume is expanded compared to the case where the totally dissimilar case.

Moving to the general case, we formalize the effect of the similarity matrix *S* as transforming the axes in the hyper-dimension space that the metacommunity is embedded into. To account for this, we simply need to compute the solid angle between all the axes. Mathematically, the solid angle Ω(*S*) (i.e., denoted with gray curves in Fig. 6) formed by the similarity matrix *S* is given by (Ribando, 2006; Song *et al*., 2018)

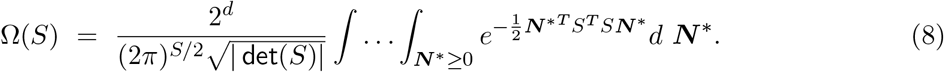

With the similarity matrix *S*, the hypervolume is transformed into Ω(*S*) vol(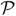). Intuitively, we can define diversity *β*_vol_ accounting for species similarity as

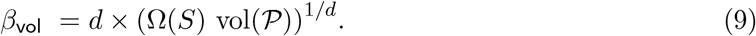

As the elements *S*_*ij*_ are always larger than 0, the transformed hypervolume and the associated beta diversity is always smaller accounting for species similarity. This aligns with ecological expectation because species being more similar would reduce the overall variation in the metacommunity.

In parallel to species similarity, we can also consider species functional complementarity. Functional complementarity means that two species provide additional ecological functioning than the addition of the functioning when both species are isolated (i.e., in monoculture) (Tilman *et al*., 2014). Multiple methods are available to quantify functional complementarity from experiments (e.g., Loreau & Hector 2001; Alahuhta *et al*. 2017). We represent functional complementarity using the *S* matrix, where *s*_*ij*_ now denotes the level of functional complementarity species *j* provides to species *i*. Note that *s*_*ij*_ are negative, as they represent functional complementarity. Because of the negative *s*_*ij*_, the hypervolume is now expanded (Figure 6D) compared to the case without any functional complementarity (Figure 6A). In general, the transformed hypervolume is always larger accounting for functional complementarity. This aligns with the ecological expectation because more variations in ecosystem functioning would increase the overall variation in the metacommunity.

While species similarity and functional complementarity are related concepts, they can have different focuses. For example, accounting for species similarity allows quantifying phylogenetic beta diversity (Graham & Fine, 2008), while accounting for functional complementarity allows quantifying functional beta diversity. As a side note, the formulas accounting for functional complementarity are identical to those for species similarity (Eqns. 8 and 9). Despite the apparent symmetry between species similarity and functional complementarity, this extension of functional complementarity is not obvious to achieve using the traditional formalism using Hill’s number in the framework of Leinster & Cobbold 2012 (but see Scheiner *et al*. (2017) for the case of functional diversity).

### 3.5 Nestedness-turnover decomposition as filling-finding facets

Decomposing beta diversity into turnover and nestedness components is a major advance in our understanding of beta diversity (Baselga, 2012; Legendre, 2014). Turnover (also known as replacement) means that species compositions tend to replace each other along spatial gradients. Nestedness (also known as richness difference) means that species composition in a community is a strict subset of the species composition in a richer community. Here, we provide a geometric interpretation of the nestedness-turnover decomposition.

For illustrative purposes, let us consider two metacommunities with one showing complete turnover and the other one being completely nested. The metacommunity matrix describing the metacommunity with complete turnover is (the corresponding geometric embedding illustrated in Figure 7A):

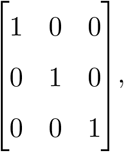

and the metacommunity matrix of the nested metacommunity is (the corresponding geometric embedding illustrated in Figure 7B):

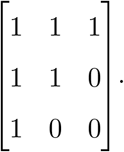

**Figure 7:**
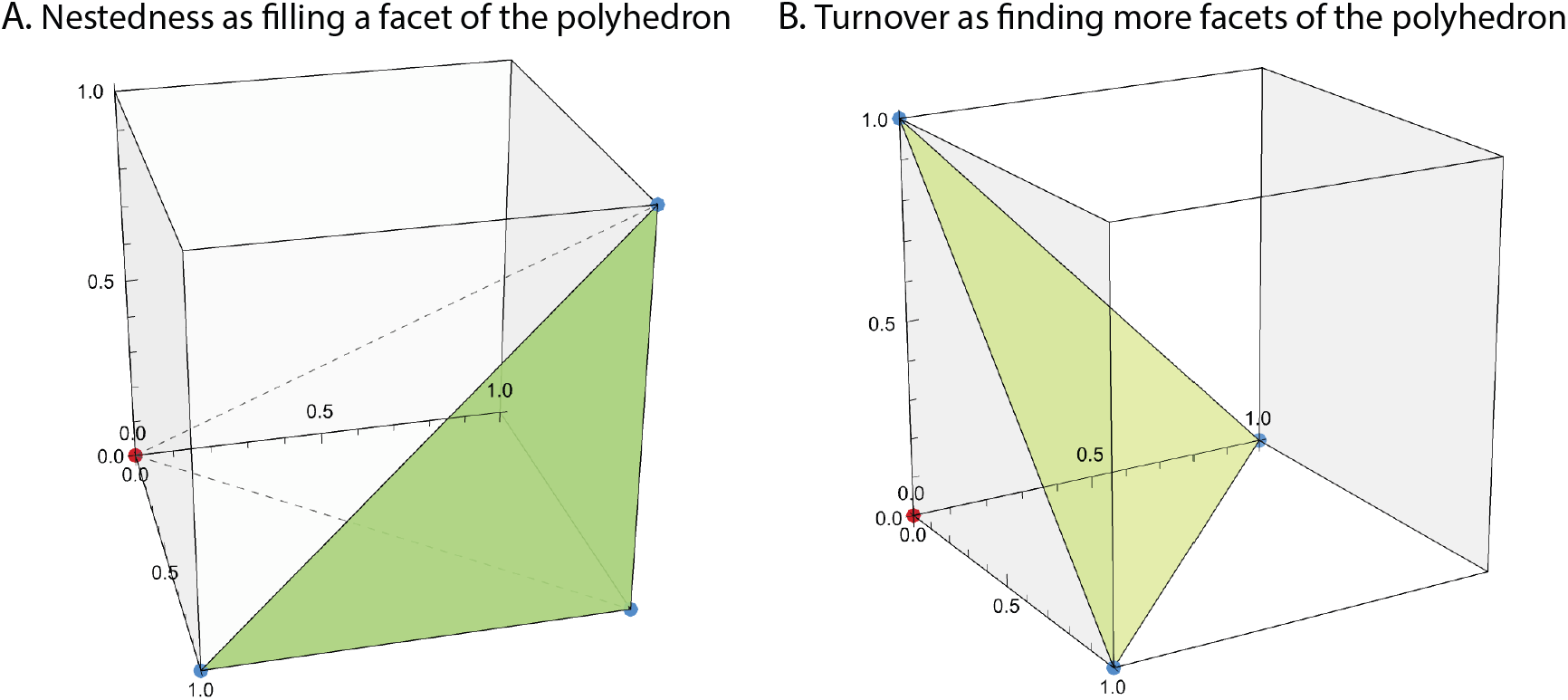
Geometric interpretation of nestedness and turnover decomposition. We consider here two metacommunities both with 3 communities and 3 species. Panel (A) shows the archetypical example of nestedness. All the points are located on the *same* facet of the 3-dimensional cube. The geometric interpretation of nestedness is to *fill* more facets of the cube. Panel (B) shows the archetypical example of turnover. All the points (each represents a community) are located on the *different* facets of the 3-dimensional cube. The geometric interpretation of turnover is to *find* more facets of the cube. These geometric interpretations generalize to higher dimension, where we replace cube as high-dimensional polyhedron.

To compare the geometric embeddings of the two metacommunities, we observe that all the embedded points are located on *different* facets of the cube in the turnover metacommunity, while all the embedded points are located on the *same* facet of the cube in the nested community. In other words, the turnover process increases the beta diversity by finding new facets, while the nestedness process increases the beta diversity by filling a facet.

In contrast to previous sections, we did not provide an analytic measure to partition geometric beta diversity into nestedness and turnover parts. This is because our geometric approach suggests that this problem may be inherently ill-defined: nestedness is essentially a multidimensional property that cannot be reduced into a single scalar index. As a metacommunity with *γ* diversity has 2*γ* facets, nestedness should be represented as a 2*γ*-dimensional vector, where each element denotes how much each facet is filled. To make it even more complicated, each element in the nestedness vector is interwinded with another element, as filling one facet can affect how another facet is filled. Thus, it is difficult, if not impossible, to summarize the nestedness vector into a 1-dimensional index without losing ecological information. Our observation complements the arguments that nestedness and turnover are interactive and thus cannot be partitioned (Šizling *et al*., 2022).

#### Box 1

**Linking the geometric approach to traditional formalisms**

Despite the differences of our geometric approach to traditional formalisms, our approach has a strong connection to them. The bridge across different formalisms exists by forming different geometric shapes from the same embedded points (communities) of the metacommunity. The main text has focused on forming a convex hull from the embedded points. However, there are other alternative choices (such as an ellipse). Different geometric shapes would result in different hypervolumes (consequently, different beta diversity). Importantly, different shapes (i.e., geometric beta diversity) emphasize different ecological properties. Here, in addition to the convex hull approach in the main text, this box introduces two other geometric approaches in forming shapes. Appendix D provides intuitions behind these two definitions, as well as mathematical derivations.

The first approach connects with the dominant formalism of beta diversity based on generalized variance (Legendre & De Cáceres, 2013). We define the hypervolume as determinant of the covariance matrix of the metacommunity matrix. The geometric interpretation of this hypervolume is the corresponding ellipse formed by the embedded metacommunity (Lu *et al*., 2021). Similar to Eqn. (4), the geometric beta diversity *β*_VAR_ is defined as *d ×* det(VAR(*X*))^1*/d*^ where *X* is the metacommunity matrix. Figure 8A-B illustrates two examples of metacommunity with *β*_VAR_. This formalism naturally partitions composition variation into a traditional beta diversity measure and a spatial association component.

**Figure 8:**
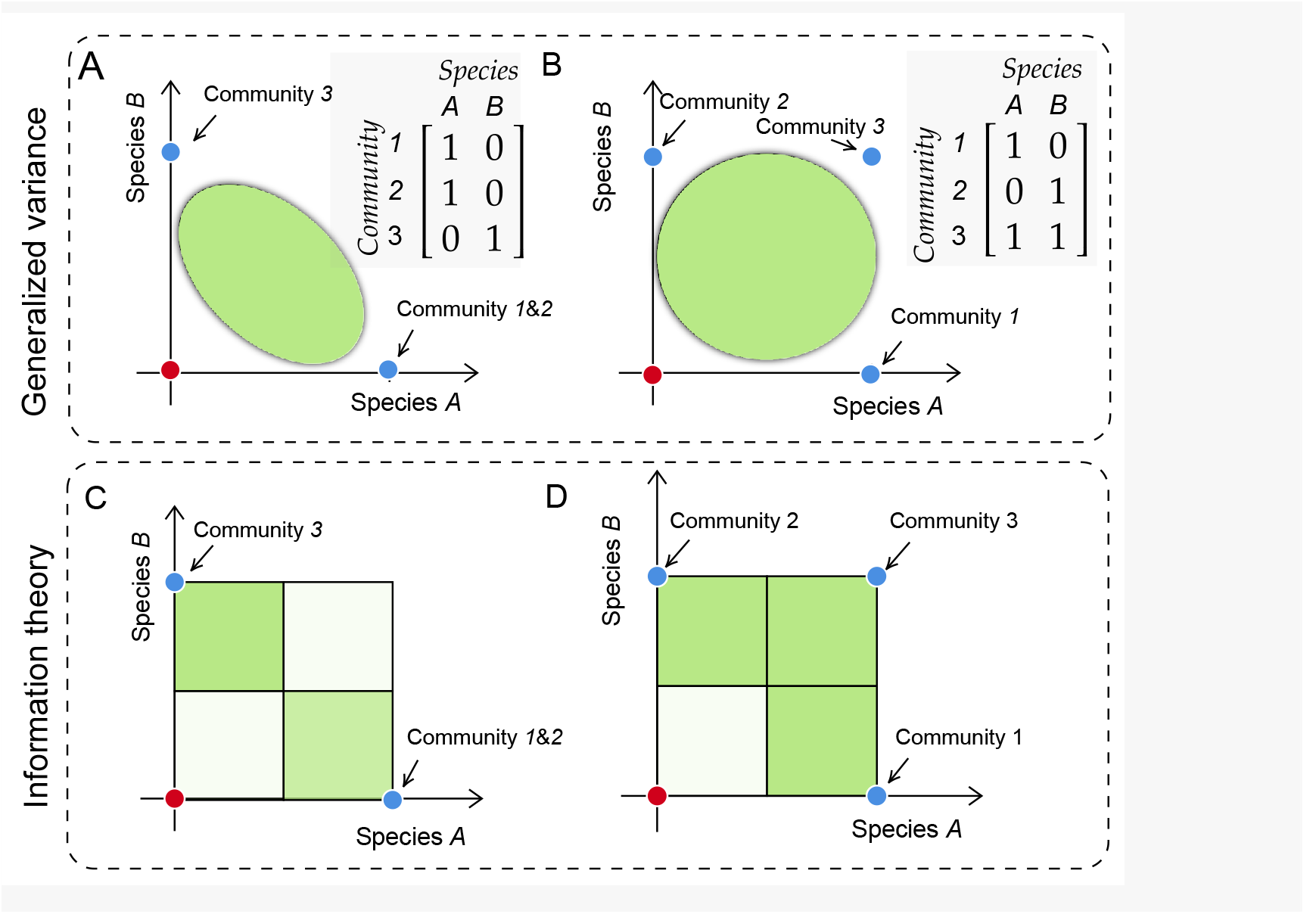
Connecting hypervolume beta diversity to traditional formalisms. We consider again metacommunity I and II from Figure 1A). Instead of forming a convex hull from the geometric embedding (Figure 1), we form either ellipse or effective support size from it. Panels (A) and (B) focus on the hypervolume of elliptic shapes formed by embedded metacommunity. This geometric shape closely connects with generalized variance. Specifically, the shape of the ellipse is determined by the structure of the covariance matrix. Panel (C) and (D) focus on the hypervolume of a multivariate Bernoulli random variable as entropy (a.k.a., the effective size of the support; Grendar 2006). Darker shading of the color represents a higher contribution of the state to the total entropy (given by the terms in Eqn. S9). The metacommunity with more evenly distributed community compositions has a higher effective size of the support and therefore higher beta diversity.

The second approach connects with the dominant formalism of beta diversity based on information theory (Chao *et al*., 2014). We define the hypervolume as the joint entropy *H*(*X*) of a multivariate Bernoulli distribution. The geometric interpretation of this hypervolume is the effective size of the support formed by the embedded metacommunity (Grendar, 2006). Similar to Eqn. (4), the geometric beta diversity *β*_info_ is defined as *d × H*(*X*)^1*/d*^ where *X* is the metacommunity matrix. Figure 8C-D illustrates two examples of metacommunity with *β*_info_. This formalism allows interpreting beta diversity in the language of information, e.g., spatial association as mutual information. This formalism is also closely connected with zeta diversity (Hui & McGeoch, 2014).

**Table 1:** Summary of geometric beta diversity metrics.

| | $\beta_{\text{vol}}$ | $\beta_{\text{VAR}}$ | $\beta_{\text{info}}$ |
| --- | --- | --- | --- |
| Geometric interpretation | Minimum convex hull volume | Ellipse volume | Effect support size |
| Community/species contribution | $\Delta\beta_{\text{vol}}$ | Univariate variance | Marginal entropy |
| Association contribution | $\Delta\beta_{\text{vol}}$ | Determinant of correlation matrix | Mutual information |
| Data type | Arbitrary measure | Arbitrary measure | presence/absence only |
| Species similarity/<br>complementarity | Both | Only similarity | No |

## 4 Empirical applications

### 4.1 Efficient estimation of beta diversity

To apply our measures of beta diversity to empirical data, we need to estimate the hypervolumes of the embedded metacommunity. The hypervolume of geometric shape in high dimension is notoriously difficult to estimate. Fortunately, we do not need to compute the hypervolume of arbitrary geometric shapes (e.g., this is typically required for fundamental niches). Appendix E discusses how to compute hypervolume beta diversity (*β*_vol_, *β*_VAR_ and *β*_info_) in detail. We have provided an R package `betavolume` (https://github.com/clsong/betavolume) to assist with these calculations. In brief, the *exact* hypervolume is only computationally feasible for metacommunities with 15 or fewer communities or species, while the *robust approximated* hypervolume is computationally feasible for metacommunities that are much larger (even for more than 10, 000 species or communities). A detailed discussion can be found in Appendix E. This package provides a user-friendly interface in R language to compute beta diversity *β*_vol_ and its various extensions (including dupli-cations in presence/absence data, community/species-specific contribution, species similarity and functional complementarity).

### 4.2 Latitudinal pattern of beta diversity

Through the years, a high-profile debate has centered on latitudinal patterns of beta diversity (Currie *et al*., 2004; Kraft *et al*., 2011; Qian *et al*., 2013; Xing & He, 2021). The dataset used in the debate is forest transect data, which contains 198 locations along a latitudinal gradient (Gentry, 1988; Janni, 2003). Each location has a plot that can be considered as a metacommunity of 10 communities. Previous research using traditional measures of beta diversity has reached contrasting conclusions: beta diversity decreases along the absolute latitude gradient when using Whittaker’s multiplicative measure (Currie *et al*., 2004), while it shows a null pattern with absolute latitude when using an alternative measure known as beta deviation (Kraft *et al*., 2011). Importantly, both patterns originate from the exponential decrease of gamma diversity along the absolute latitude gradient (Figure 9C). Specifically, the pattern with Whittaker’s multiplicative measure is fully driven by gamma diversity, as an exponential decrease in gamma diversity completely masks the effects of alpha diversity. In contrast, the pattern with beta deviation is due to the ignorance of gamma diversity, as beta deviation removes the effect of changing gamma diversity (Bennett & Gilbert, 2016). However, both metrics ignore the interactive effect of nestedness and turnover in shaping the latitudinal pattern of beta diversity.

**Figure 9:**
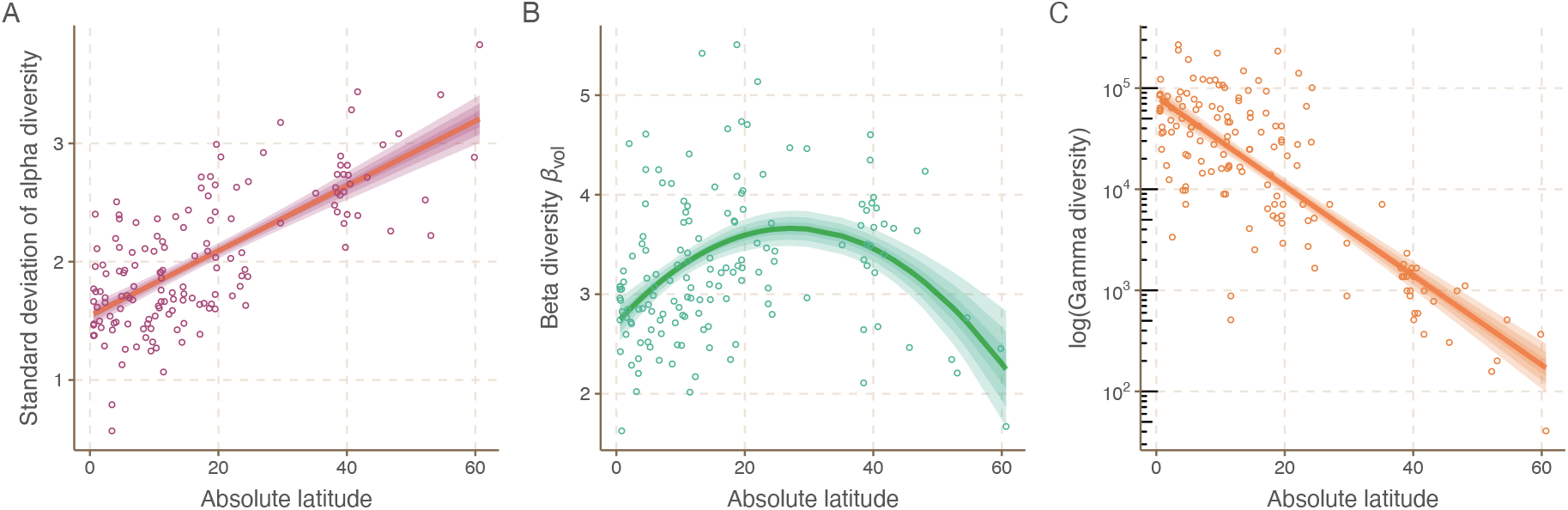
The pattern of beta diversity *β*_vol_ along latitudes and its origin. We show how (variance of) alpha diversity (Panel A), beta diversity (Panel B), and gamma diversity (Panel C) changes along the absolute latitude gradient. The horizontal axis shows the absolute latitude, while the vertical axis shows the measure of diversity. Each point represents a metacommunity. We depicted the generalized additive line with shaded confidence intervals. Panel (A) uncovers a monotonically increasing trend of the variance of alpha diversity (adjusted *R*^2^ = 0.64). Panel (B) shows our measure *β*_vol_ has a unimodal pattern (*p* = 0.99 according to Hartigans’ dip test; Hartigan & Hartigan 1985). This is in direct contrast to previous results, where beta diversity is either monotonically decreasing or does not change. Panel (C) shows gamma diversity exponentially decreases (adjusted *R*^2^ = 0.65).

We applied our measure *β*_vol_ to this dataset (Gentry, 1988; Janni, 2003). In contrast to the previous consensus, we find a unimodal pattern of *β*_vol_: it first increases and then decreases along the absolute latitude gradient (Figure 9B). This pattern emerges from the conflicting trends of the nestedness and turnover components of beta diversity. On the one hand, the decreasing gamma diversity has a negative effect on the turnover component of the *β*_vol_ (Figure 9C), because loosing a species in the regional species pool is equivalent to loosing a facet in the multivariate geometric space (see Figure 7B in section 3.5). On the other hand, the increasing variance in alpha diversity has a positive force on the nestedness component of the *β*_vol_ (Figure 9A). This is because the increasing difference in alpha diversity across communities increases the chance of filling a facet (Figure 7A). The high gamma diversity and low variance of alpha diversity in the lower latitude suggest that the metacommunities in the region are characterized by strong mutual exclusions among species, while the low gamma diversity and high variance of alpha diversity in the higher latitude suggest that the metacommunities are characterized by high nestedness. The unimodal pattern of *β*_vol_ therefore indicates that the mid-latitude metacommunities have the richest community types. The unimodal pattern and the peak at around 30 degree absolute latitude are consistent with observations from another global dataset on plant diversity (Scheiner & Rey-Benayas, 1994).

Note that the unimodal pattern is not our key take-away. Given the spatial and temporal biases in global biodiversity datasets (Gonzalez *et al*., 2016; Hughes *et al*., 2021), there is plenty of room for disagreement on which is the true latitudinal pattern of beta diversity. Nonetheless, as the nestedness and turnover components interactively shape beta diversity gradients, a satisfactory measure of beta diversity should be able to account for the effect of both. Our measure *β*_vol_ is capable of doing this, while previous measures either mostly only extract the information captured by gamma diversity or inappropriately account for the effect of the nestendess and turnover components (Šizling *et al*., 2022).

### 4.3 How sampling efforts affect beta diversity

In empirical estimation of beta diversity, sampling efforts play a prominent role. That is, with traditional measures of multiple-site beta diversity, beta diversity always increases when more sites are sampled. This increase in beta diversity is mainly driven by the increase in gamma diversity (Bennett & Gilbert, 2016; Xing & He, 2021). However, this begets two problems: first, more sampling may not pay off, as it provides exponentially diminishing returns; second, we cannot distinguish which metacommunity is more spatially heterogeneous. A potential solution to these problems is to implement some scaling method to adjust traditional measures according to sampling effort, ensuring a more standardized comparison. Nevertheless, our measure *β*_vol_ inherently addresses these problems without necessitating any modifications. Unlike traditional measures, *β*_vol_ does not automatically increase with greater gamma diversity. Instead, an increase in gamma diversity expands the dimensionality of the metacommunity’s embedded space, which could lead to a decrease in the rescaled hypervolume. This characteristic of *β*_vol_ allows a more accurate measure of spatial heterogeneity of metacommunities without being disproportionately influenced by the number of species (gamma diversity) or the scale of sampling efforts.

As a proof of concept, we focused on two datasets from Bennett & Gilbert (2016). One dataset contains 1-m^2^ plots in early successional fields in the Koffler scientific reserve in Ontario, Canada. Another dataset contains 50-m^2^ forest plots at Mont St. Hilaire near Montreal, Canada (Gilbert & Lechowicz, 2004). These two datasets were collected for different purposes. The data from Koffler Scientific Reserve were designed to sample a relatively homogeneous area, while the data from the Mont St. Hilaire were acquired to capture environmental heterogeneity. Previous research has shown that traditional beta diversity in both datasets would increase with sampling effort with a power-law scaling (Bennett & Gilbert, 2016; Xing & He, 2021). Thus, traditional measures fail to capture the ecological differences between the two datasets.

We apply our measure *β*_vol_ to these two datasets (Figure 10) using the random subsampling procedure of Bennett & Gilbert (2016). At Mont St. Hilaire, *β*_vol_ consistently increased with sampling effort, aligning with the expectation that more intensive sampling in a heterogeneous environment uncovers greater species turnover between plots. Conversely, at the Koffler Scientific Reserve, *β*_vol_ initially increased but then declined and plateaued. This pattern is consistent with sampling in a more homogeneous environment, where the majority of species turnover is captured at lower sampling efforts, and additional sampling yields diminishing returns in terms of detecting new compositional differences.

**Figure 10:**
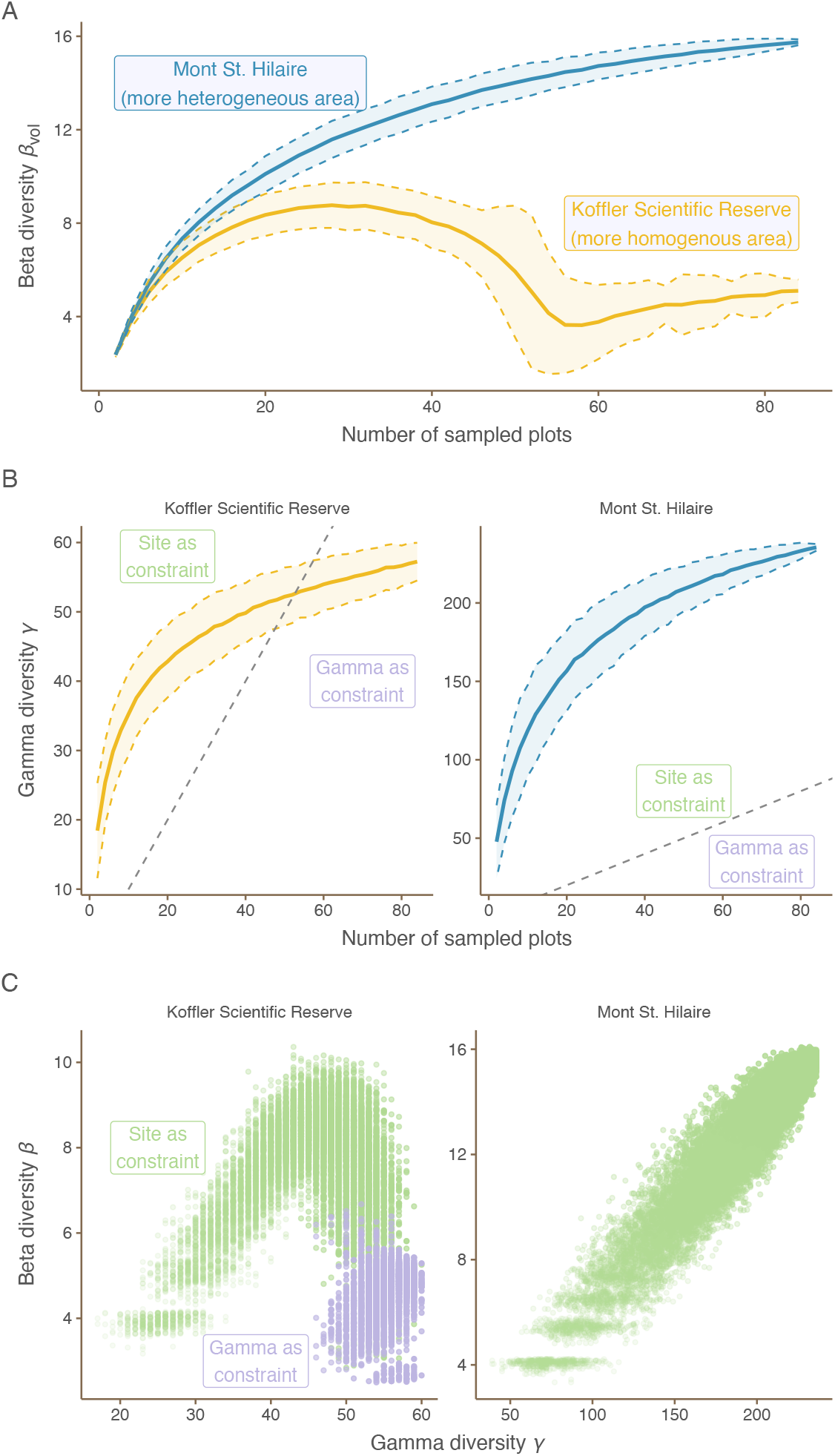
Relationship between sampling effort and biodiversity across two sites: the Koffler Scientific Reserve (a more homogeneous area) and Mont St. Hilaire (a more heterogeneous area). Panel (A): Beta diversity (*β*_vol_) increases with sampling effort at Mont St. Hilaire (blue), while it plateaus at the Koffler Scientific (orange) Reserve after an initial increase. This indicates that Mont St. Hilaire harbors higher species turnover between plots than the Koffler Scientific Reserve. Shaded areas represent two standard deviations around the mean. Panel (B): Gamma diversity (*γ*) increases with sampling effort at both sites, following a power-law relationship. This aligns with the well-known species-area relationship. The constraint on beta diversity calculation (dimension of embedding) is determined by the minimum of gamma diversity (purple) and the number of communities (green). For Mont St. Hilaire, the site is always the limiting factor, whereas for the Koffler Scientific Reserve, the constraint shifts from site to gamma diversity with increased sampling effort. Panel (C): The relationship between beta diversity *β*_vol_ and gamma diversity *γ* reveals that beta diversity at the Koffler Scientific Reserve declines before the constraint switches from site to gamma diversity (green to purple). This suggests that the decrease in beta diversity reflects a true pattern of homogeneity rather than an artifact.

Interestingly, the decline in *β*_vol_ at the Koffler Scientific Reserve occurs before the constraint on the embedding dimension switches from site to gamma diversity (Figure 10B and C). This suggests that the plateau in *β*_vol_ reflects a genuine ecological pattern of homogeneity rather than an artifact of sampling limitations. Thus, our measure *β*_vol_ has the potential to solve the long-standing issue on sampling efforts associated with beta diversity: more sampling is necessary to detect the ecological differences in spatial heterogeneity between the two datasets.

## 5 Discussion

Beta diversity is a central concept in spatial ecology and conservation management (Mori *et al*., 2018). Unlike the consensus on measures of alpha and gamma diversity (at least for presence/absence data) (Jost, 2007; Chao *et al*., 2014), there is a long list of beta diversity measures. One may argue that the pressing problem now should be classifying or reconciling these measures of beta diversity (Jurasinski *et al*., 2009). If this is so, then why introduce a new measure at this point? We believe our new measure of beta diversity is much needed because (1) it reconceptualizes beta diversity, (2) its geometric nature makes it easily extendible and generalizable, (3) it synthesizes traditional measures, and (4) it provides novel ecological insights. We discuss these four advantages below.

First, our new measure behaves qualitatively differently from all traditional measures. Our measure is built upon a core observation that beta diversity should be maximized when we observe all possible community compositions in the region (Figure 1). In short, the more the merrier. In contrast, traditional wisdom posits beta diversity is maximized when each community only has a single distinct species (see review in Legendre & De Cáceres 2013). The traditional wisdom operates under an individualist perspective. That is, the ‘ecology’ of a species is the same with or without the presence of other species. Under this individualist perspective, each community with a single unique species would display the maximum possible variance of biodiversity in the region. However, the individualist perspective is unlikely to be general in ecological systems where interactions abound. Thus, our measure is conceptually justified as long as species interactions in a local community affect species dynamics and functioning.

It is not a trivial problem to formalize this reconceptualization of beta diversity. To our knowledge, among the traditional measures, the only exception to traditional wisdom is the Shannon diversity of realized species combinations (Juhász-Nagy & Podani, 1983). This measure proposed to simply count the number of unique community compositions (Juhász-Nagy & Podani, 1983). However, this measure ignores quantitatively how community compositions are different. For example, a community with species *A* and *B* should be more distinct from a community with species *C* than a community with composition *A*. We have taken a hypervolume approach to solve this problem. Hypervolume is an old friend in ecology, and was used most famously by Hutchinson to frame the discussion of the niche (Blonder, 2018). The idea of hypervolume has been widely used in various areas of ecology research (Raup & Michelson, 1965; Violle & Jiang, 2009; Boucher *et al*., 2013; Blonder *et al*., 2014). Notably, researchers have measured functional beta diversity as the *overlap* between the functional trait spaces of two local communities (Mammola, 2019; Lu *et al*., 2021). In contrast to these previous works, our measure is fundamentally different, as we directly interpret hypervolume of the metacommunity matrix as beta diversity. To do so, we have followed the idea of Hutchinson where he interpreted the fundamental niche as hyper-dimensional geometric shapes (Hutchinson, 1957). Our geometric measure provides a linear scaling between beta diversity and the number of unique community compositions, while it also quantifies the difference between unique community compositions (Figures 2 and S3).

Second, our approach provides a unifying framework for beta diversity. Given the importance of beta diversity, the basic quantification is far from enough for empirical study. We have extended our geometric measure to the following five cases with strong empirical importance: duplications in presence/absence data (Figure 3), temporal changes (Figure 4), community/species-specific contributions to beta diversity (Figure 5), species similarity and functional complementarity (Figure 6), and turnover-nestedness decomposition (Figure 7). While these extensions are possible with traditional measures of beta diversity, they often, although not always, require different theoretical formalisms. In part, this may result from the fact that most traditional measures are algebraic manipulations of metacommunity matrix without a simple geometric interpretation. In contrast, we present a geometric approach, which is fully visual in 2- or 3-dimensional space. This visual aspect of our geometric approach permits an intuitive and generalizable ecological interpretation. A psychological benefit with our approach is that humans are intrinsically more familiar with geometry than algebra (Sablé-Meyer *et al*., 2021). Thus, our geometric measure is, in general, easier to visualize, interpret, and generalize than traditional algebraic definitions.

Third, our measure provides a unifying approach to synthesize previous measures of beta diversity. We are not simply adding yet another measure to the list of beta diversity measures. Instead, our measure considers new higher-order information that traditional measures have missed. Despite the variety of traditional measures, most of them can be classified into two schools of thought: variance-based or information-based. The variance-based measure considers the diagonal of the covariance matrix (Legendre & De Cáceres, 2013), while we have in addition considered the off-diagonal. These off-diagonal components represent ecologically the spatial associations of species (Figures 1 and 8). The most commonly used information-based measure considers the pooled marginal entropy of a joint distribution (Jost, 2007), while the joint entropy takes the mutual information into account (Juhász-Nagy & Podani, 1983). In other words, previous measures of beta diversity have a geometric basis, and our approach reveals their hidden geometric nature.

Fourth, our measure provides novel ecological insights into the patterns in empirical data. We have focused on two important empirical issues: global syntheses of biodiversity data and the sampling efforts. Focusing on global syntheses, traditional measures are masked by the exponentially changing gamma diversity; thus, the latitudinal pattern is mostly driven by gamma diversity. In contrast, our measure can reveal the joint effects of alpha and gamma diversity in shaping the patterns of beta diversity (Figure 9). Focusing on the sampling efforts, traditional measures fail to reveal additional information with increasing sampling effort. This is because traditional measures are again masked by increasing gamma diversity with increasing sampling effort. In contrast, we show that increasing sampling effort is necessary to detect hidden spatial heterogeneity, and our measure can help quantify this heterogeneity (Figure 10). Besides the demonstrated examples, we also expect that our metric should be particular useful in determining the relationship between species composition and ecosystem functioning (Grman *et al*., 2018; Mori *et al*., 2018) and stability (McGranahan *et al*., 2018) because it explicitly takes species-association into account. When applied in the temporal context, the hypervolume-based beta diversity is also a measure of community change predictability (Song *et al*., 2021; De Cáceres *et al*., 2019): for example, in time-lag analysis, higher beta diversity indicates more random community composition changes over time while lower beta diversity indicates more directional changes (Jones *et al*., 2017).

We want to emphasize that our geometric approach is not meant to replace existing measures but rather to provide a complementary perspective. For example, in conservation policy-making, traditional measures may be more suitable, as they focus on identifying areas with unique species compositions or those that contribute significantly to regional diversity. Nevertheless, when the goal is to understand the ecological processes underlying biodiversity patterns or to assess the functional consequences of biodiversity loss, our geometric approach can provide insights by capturing the non-additivity caused by the complex interactions among species. In sum, as no single metric can encompass the full complexity of biodiversity, researchers should carefully select the beta diversity measure that best aligns with their specific research questions and the ecological context of their study.

Like other beta diversity metrics, our method is not without limitations. One major issue is that hypervolume beta diversity is sensitive to normalization of the elements in the metacommunity matrix (Legendre & De Cáceres, 2013). For example, beta diversity is likely to be different when we consider the absolute versus relative species abundance. However, we consider it to be a feature rather than a bug. For example, under some ecological rationales, we can argue that a metacommunity with more individuals, *ceteris paribus*, is more “diverse” than another metacommunity with fewer individuals (Legendre & De Cáceres, 2013). We suggest that every normalization method requires careful ecological interpretation. As long as we apply the same normalization method across metacommunities of interests, we can safely compare which metacommunity has a higher beta diversity (in accordance with the ecological rationale behind the normalization).

We believe that our proposed measure is readily applicable to existing data. To further expand its applicability, we envision the following extensions of our geometric framework. One future direction is to further explore geometric features of the embedded metacommunity. For example, we have not yet considered its geometric asymmetry. For example, let us consider two metacommunities are both embedded as triangles with identical volume, but one is equilateral while the other one is not. The ecological differences between them is that the equilateral metacommunity has more balanced species distributions across local communities. To quantify the association between geometric asymmetry and species balance, it might be useful to adapt tools from studies on the geometric asymmetry in different ecological contexts (Grilli *et al*., 2017; Medeiros *et al*., 2021). Another future direction is to develop analytic models of null models. Null models are widely used in beta diversity analysis to disentangle confounding factors. Analytic null models are available for many traditional measures of beta diversity (Xing & He, 2021; Lu *et al*., 2019; Lu, 2021; Deane *et al*., 2022). An analytic expression for *β*_vol_ is challenging because of the complexity in quantifying hypervolume (Appendix E). However, *β*_VAR_ and *β*_info_ are tightly linked to high-dimensional normal distributions, thus it is possible to obtain analytic expressions. Furthermore, a promising future direction is to extend our geometric approach to measure functional diversity (reviewed in Scheiner *et al*. 2017).

## 6 Conclusion

We have reconceptualized the measurement of beta diversity as a geometric hypervolume. We have shown the connections of our new measure to existing variance- and information-based measures. Our geometric approach provides a unified way to measure beta diversity that can deal with duplications in presence/absence data, temporal change, turnover, nestedness, species and functional complementarity. We demonstrated its application to two datasets and the novel insights it offers. We have provided the tools needed to apply our approach. This new framework has the potential to deepen our understanding of biodiversity patterns and processes across scales by explicitly considering the complex interactions and non-additivity inherent in ecological systems.

